# Influenza A virus diffusion through mucus gel networks

**DOI:** 10.1101/2020.08.14.251132

**Authors:** Logan Kaler, Ethan Iverson, Shahed Bader, Daniel Song, Margaret A. Scull, Gregg A. Duncan

**Author notes:** Correspondence to: Gregg Duncan.

## Abstract

In this study, influenza A virus (IAV) and nanoparticle diffusion in human airway mucus was quantified using fluorescent video microscopy and multiple particle tracking. In previous work, it was determined that mucin-associated sialic acid acts as a decoy receptor for IAV hemagglutinin binding, and that virus passage through the mucus gel layer is facilitated through the sialic-acid cleaving enzyme, neuraminidase (NA), also present on the IAV envelope. However, our data suggests the mobility of IAV in mucus is significantly influenced by the mesh structure of the gel, as measured by nanoparticle probes, and NA activity is not required to facilitate virus passage through mucus gels. Using newly developed analyses, the binding affinity of IAV to the 3D mucus meshwork was estimated for individual virions with dissociation constants in the mM range, indicative of weak and reversible IAV-mucus interactions. We also found IAV diffusion significantly increased in mucus when treated with a mucolytic agent to break mucin-mucin disulfide bonds. In addition, IAV diffusion was significantly limited in a synthetic mucus model as crosslink density was systematically increased and network pore size was reduced. The results of this work provide important insights on how the balance of adhesive and physical barrier properties of mucus influence the dissemination of IAV within the lung microenvironment.

## INTRODUCTION

Influenza virus has the ability to rapidly evolve, quickly spreads, and poses a major health threat with a high morbidity and mortality rate (1, 2). Individuals with chronic lung disease, including asthma and chronic obstructive pulmonary disease (COPD), have an increased risk of potentially life-threating medical complications due to influenza virus infection (3, 4) To protect the respiratory tract from infection by influenza virus and other pathogens, a thin layer of mucus is constantly produced and coats the surface of the airway epithelium (5). Mucus is a hydrogel composed of approximately 98% water and 2% solids, containing secreted mucins, globular proteins, lipid surfactants and cellular debris (6, 7). The primary building blocks of mucus gels are high molecular weight (≥ 1 MDa) mucin biopolymers that are heavily glycosylated with sugars such as fucose, sulfate, and sialic acid (7–9). Mucin biopolymers interweave to form a gel network with a pore size ranging from 100-500 nm, depending on total mucin concentration (8). Ciliated cells on the airway epithelium beat in a coordinated fashion to clear mucus and trapped particulates from the airway (10). In order to prevent infection, mucus in the lung must effectively trap inhaled pathogens and the mechanisms by which this is achieved are important to our understanding of innate host defenses.

To initiate infection, the influenza virus lipid envelope contains receptor-engaging hemagglutinin (HA) glycoproteins and receptor-destroying neuraminidase (NA) enzymes (11). HA binds to α-2,3 and/or α-2,6 linked sialic acid on the surface of the cell to allow for internalization of the virus (12, 13). The various strains of influenza virus have different sialic acid binding preference, with human viruses typically preferring α-2,6 linked sialic acid while avian and equine viruses prefer α-2,3 linked sialic acid (14). Newly formed virions are released from the cell when NA catalyzes the cleavage of terminal sialic acid (15). Previous work has shown the same HA/NA machinery also plays a role in the penetration of IAV through the mucus barrier to infection (13, 16, 17). Specifically, prior work has shown HA on IAV binds to sialic acid within mucus, which is predominantly as α-2,3 linked sialic acid on mucins (18). In order to facilitate IAV diffusion through mucus, NA activity is required for removal of sialic acid decoy receptors (17). However, whether this can be generalized to human influenza virus infections remains unclear as the evidence to support this mechanism is based on data generated using non-human IAV, non-human mucus, and synthetic glycan-coated substrates. In addition, prior work on nanoparticle and virus diffusion has noted substantial variation between the barrier properties of human mucus between individual donors (19–21) which would likely contribute to differences in their susceptibility to infection. Currently, it is unknown if factors, such as mucus gel pore size and/or mucin glycan profile, influence host-specific mucus barrier properties towards IAV and other respiratory viruses.

In this work, we use fluorescent microscopy to measure diffusion of IAV through human mucus. Our investigations were conducted using the well-characterized H1N1 influenza A/Puerto Rico/8/34 (PR8) strain, of human origin but with an ‘avian-like’ HA binding preference for α-2,3 linked sialic acid (22). We collected human samples from 10 patient donors for our studies to assess variation in their barrier properties. An important component of our work was also measuring diffusion of nanoparticles designed to be minimally adhesive to mucus (19, 23) with a comparable size to IAV (∼100 nm) in each sample tested. This enabled us to develop an analytical model to decouple physical and adhesive barrier functions of mucus towards IAV. This technique provides a method to assess effective dissociation constants for IAV and potentially other respiratory viruses in 3D human mucus gels. The results of this work give new insight into how the structural and adhesive properties of mucus influence its protective function against IAV infection.

## MATERIALS AND METHODS

### IAV and Nanoparticle Preparation

The plasmid-based reverse genetics system for influenza virus A/Puerto Rico/8/34 (PR8; H1N1) was a gift from Adolfo Garcia-Sastre. Infectious virus was generated from cloned cDNAs in 293T and Madin-Darby canine kidney (MDCK) cell co-cultures and purified on a sucrose cushion as previously described (24, 25). Viral infectivity of resulting virus stocks was quantified by standard plaque assay on MDCK cells (26), yielding 3.9×10^9^ plaque-forming units (pfu)/mL. IAV was subsequently labeled with a lipophilic dye, 1,1’-dioctadecyl-3,3,3’3’-tetramethylindocarbocyanine perchlorate (DiI; Invitrogen). The labeled virions were then purified and concentrated via haemadsorption to chicken red blood cells (27). IAV stock was aliquoted and stored at -80°C. Each aliquot underwent a maximum of two freeze thaws. This strain exhibits a primarily spherical morphology and diameter of roughly 100 nm. DiI-labeled virus stocks were counter-stained with polyclonal anti-influenza virus H1 (H0) HA PR8 antibody (NR-3148; antiserum, goat; BEI Resources, NIAID, NIH). Carboxylate modified fluorescent PS nanoparticles (PS-COOH; Life Technologies) with a diameter of 100 nm were coated with a high surface density of polyethylene glycol (PEG) via a carboxyl-amine linkage using 5-kDa methoxy PEG-amine (Creative PEGWorks) as previously reported (19). The NanoBrook Omni (Brookhaven Instruments) was used to conduct dynamic light scattering (DLS) experiments to determine virus and nanoparticle size distribution.

### Neuraminidase Assay

Neuraminidase activity was tested using the NA-Fluor™ Neuraminidase Assay Kit from Life Technologies and following manufacturer instructions. Fluorescence intensity was measured using a Spark Multimode Plate Reader (Tecan). A standard curve for 4-Methylumbelliferone (4-MU(SS); Sigma-Aldrich) was then generated and the fluorescence signal of 18,000 relative fluorescent units (RFU) corresponding to 20µM 4-MU(SS) was chosen to normalize NA activity. The neuraminidase assay kit was also used to perform a neuraminidase inhibition assay using the neuraminidase inhibitor zanamivir (Cayman Chemical). IAV neuraminidase inhibition was tested using a 1:16 dilution of IAV and varying the concentration of zanamivir from 0.01 nM to 10 µM. In accordance with manufacturer instructions, IAV was incubated with zanamavir for 30 min before the NA-Fluor™ substrate was added. The reaction was incubated for 1 h at 37°C before fluorescence intensity was measured.

### Synthetic Mucus Hydrogel Preparation

Using a previously established method (28), synthetic mucus hydrogels were prepared using 2% porcine gastric mucin (PGM; Sigma-Aldrich) and varying percentages (1.0%, 1.5%, and 2.0%) of 4-arm polyethylene glycol (PEG; Laysan Bio Inc.) used as a thiosulfate crosslinker. The mucin and the crosslinker were combined in a physiological buffer (154 mM NaCl, 3 mM CaCl_2_, and 15 mM NaH_2_PO_4_ at pH 7.4) and mixed for 2 hours. Cross-linking solutions were prepared separately in buffer directly before mixing with mucin solutions. To initiate gelation, equal volume aliquots of each solution were mixed and equilibrated for 21 hours at room temperature. After synthetic mucus gels were fully formed, DiI-labeled IAV and/or nanoparticles were added prior to fluorescent video microscopy experiments.

### Human mucus collection

Human mucus was collected under an IRB-approved protocol at the University of Maryland Medical Center (UMMC; Protocol #: HP-00080047). Samples were collected by the endotracheal tube (ET) method, as previously described (19). ET were collected from patients after intubation as a part of general anesthesia at UMMC. In order to collect mucus from ET, the last approximately 10 cm of the tubes were cut, including the balloon cuff, and placed in a 50-mL centrifuge tube. The ET tube was suspended in the tube with a syringe needle and was then spun at 220 g for 30 seconds, yielding 100–300 μL of mucus. Mucus with visible blood contamination was not included in the analysis. Samples were stored at 80°C immediately after collection and thawed (up to 3x) prior to use for experiments. Notably, control experiments showed the microrheology of an ET mucus sample was minimally impacted by freeze-thaw as compared to an unfrozen sample kept at 4°C (**Fig. S1**).

### Fluorescent video microscopy

Samples were prepared for imaging by placing a vacuum grease coated O-ring on microscope cover glass. The sample was then applied to the center of the well and then sealed with a coverslip. Samples were allowed to stabilize for 30 minutes at room temperature prior to imaging. Slides were imaged using Zeiss LSM 800 inverted microscope with a 63x water-immersion objective. For each sample, 10 s videos were recorded at 33.3 frames per second. For all experiments, 1 µL of DiI-labeled IAV (3×10^9^ pfu/mL) and 1 µL of PEG-coated nanoparticles were added to 20 µL thawed and DiI labeled human or synthetic mucus in the center of the slide well and stirred with a pipette tip prior to imaging. After equilibration and initial imaging, samples were incubated for 15 minutes at 37°C and then imaged. For IAV with neuraminidase inhibitor (NAI) in human mucus, 1 µL of IAV labeled with DiI was mixed with 1µL of NAI Zanamivir (Cayman Chemical) and allowed to equilibrate for 10 minutes. The IAV and NAI mixture was then combined with 2.6 pM 100nm PEG-coated polystyrene nanoparticles (Invitrogen) and added to 20 µL of human mucus. The final concentration of NAI was 10 µM. For freeze-thaw testing, IAV was added to 20 µL of different aliquots of the same human mucus sample after 0, 1, 2, and 3 freeze-thaw cycles. For mucus treated with N-acetyl-L-cysteine (NAC; Sigma-Aldrich), 1 µL NAC was added to the final concentration of 50 mM.

### Multiple particle tracking (MPT) analysis

Acquired videos were processed using an in-house MATLAB (The MathWorks, Natick, MA) analysis code to isolate and track imaged particles. For each video, the mean squared displacement (MSD) was calculated as ⟨MSD(τ)⟩ = ⟨(*x*^2^+*y*^2^)⟩ (6), for each particle. The MSD values of the particles were then filtered using based on the α value, calculated as α = (*dlog*MSD(τ))/(*dlog*(τ)) (6). Particles that have an α < 0 are removed and particles with 0 < α ≤ 1 are kept, as an α = 1 as this indicative of Brownian motion (6). The alpha-filtered MSD values were then used to calculate the microrheological properties of the samples tested using the Stokes-Einstein relation (29). The equation *G*(*s*)= 2*k*_*B*_*T*/(π*as*⟨Δ*r*^2^(*s*)⟩) gives the viscoelastic spectrum where *k*_*B*_*T* is the thermal energy, *a* is the radius, and *s* is the complex Laplace frequency (19). The complex modulus is calculated by *G*^*\**^(*ω*)=*G*^*’*^(*ω*)+*G*^*”*^(*iω*) where *iω* is used in place of *s, i* is a complex number, and *ω* is the frequency (19). From this equation, the pore size (*ξ*) can be estimated as ξ = (*k*_*B*_*T*/*G’*)^1/3^ and the complex microviscosity (*η\**) can be calculated as *η*^*^= (*G*^*^(*ω*))*(*ω*) (19).

### Potential Energy and Dissociation Analysis

The trajectories of each individual particle from MPT were re-centered, based on the average *x* and *y* values, and the radial distance from the center of the trajectory was calculated for each frame of the trajectory. The radial trajectories were then put into a histogram and normalized. The normalized histogram counts were used to calculate the potential energy profile using the equation *U*/*k*_*B*_*T*= -ln[*n*(*r*)/*n*_*max*_], where *n*(*r*) is the number of counts in the normalized histogram for a given *r* value, and *n*_*max*_ is the largest count value (30). The average energy profile calculated from the polystyrene nanoparticle (PS-NP) data was subtracted from each individual profile IAV profile for a given patient sample. The harmonic well depth (*U*_M_) was calculated by determining the two maximum energy values (nonzero) for an individual particle and averaging them together. The well range (δ) was then calculated by summing the absolute values of the first and last *r* value and dividing by 2. Further calculations were not performed on *U*_*M*_ values less than 1 *kT* as this would result in a negligible IAV-mucus net interaction. The spring constant (*k*_*s*_) was calculated by *k*_*s*_=(−2*U*_*M*_/δ^2^)*r*^2^, where *U*_*M*_ is the harmonic well depth (in pN·µm), *δ* is the well range (in µm), and *r* is the displacement from the trajectory center (in µm) (31). A generalized expression was used to estimate the dissociation constant (*K*_*d*_) for fitted harmonic well potentials calculated as *K*_*d*_=(2π*k*_*B*_*T*/*k*_*s*_)^-3^/^2^ exp(-*U*_*M*_), where *U*_*M*_ is the harmonic well depth (in units of *kT*), *k*_*s*_ is the spring constant (in pN/nm), and *k*_*B*_*T* is energy (in pN·nm) (31).

### Statistical Analysis

Data was graphed and statistically analyzed using GraphPad Prism 8 (GraphPad Software, San Diego, CA).

## RESULTS

### IAV diffusivity in human mucus

To visualize IAV in human mucus, we used the lipophilic dye, DiI, to label individual virions. To confirm the identity of fluorescently-tagged particles, DiI-labeled IAV were stained with an antibody specific to IAV HA and imaged using fluorescent microscopy, revealing co-localization of the dye and viral antigen (**Fig. 1A**). Nanoparticles were included in the imaging to establish plane of view and co-staining with anti-HA antibody was not observed. We next used DLS to compare the size of unlabeled and DiI-labeled IAV. Our results indicate both labeled and unlabeled IAV were ∼120 nm in diameter, suggesting that DiI intercalation did not disrupt virions or cause particle aggregation (**Fig. 1B**).

**Figure 1.**
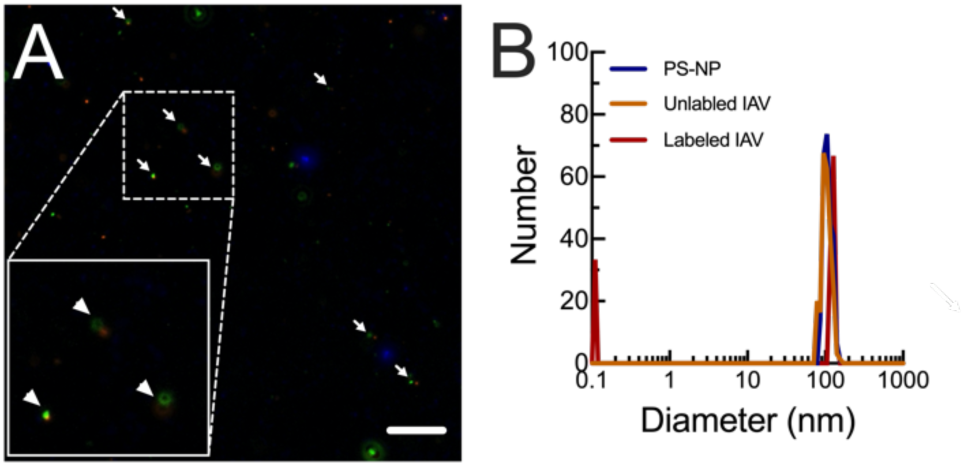
Characterization of fluorescent IAV. **(A)** Fluorescent micrograph of purified IAV and 100 nm PS-NP (blue) stained with DiI (orange) and anti-HA antibody (green). DiI and anti-HA co-staining is denoted with white arrows. Scale bar = 10 µm. **(B)** Measured hydrodynamic diameter for muco-inert PS-NP, unlabeled IAV, and DiI-labeled IAV.

After confirming IAV labeling, we introduced IAV and muco-inert PS-NP to human airway mucus and measured their diffusion using fluorescent video microscopy and MPT analysis. By simultaneously measuring PS-NP and IAV diffusion, we were able to analyze mucus microrheology and determine the diffusion rate of IAV particles within the same regions of interest. The diffusivity, as measured by logarithm based 10 of MSD at a time lag of 1 second (log_10_[MSD_1s_]), showed significant differences between PS-NP and IAV diffusion (**Fig. 2A,B**). The MSD of the IAV was, on average, 3-fold lower than that of the PS-NP (**Fig. 2C**). In addition, measured MSD for IAV varied over several orders of magnitude between individual patient samples. When comparing microrheology, as measured by PS-NP, and IAV diffusion from individual patients (**Fig. 2D**), we found mucus gel pore size positively correlates with the diffusivity of IAV particles.

**Figure 2.**
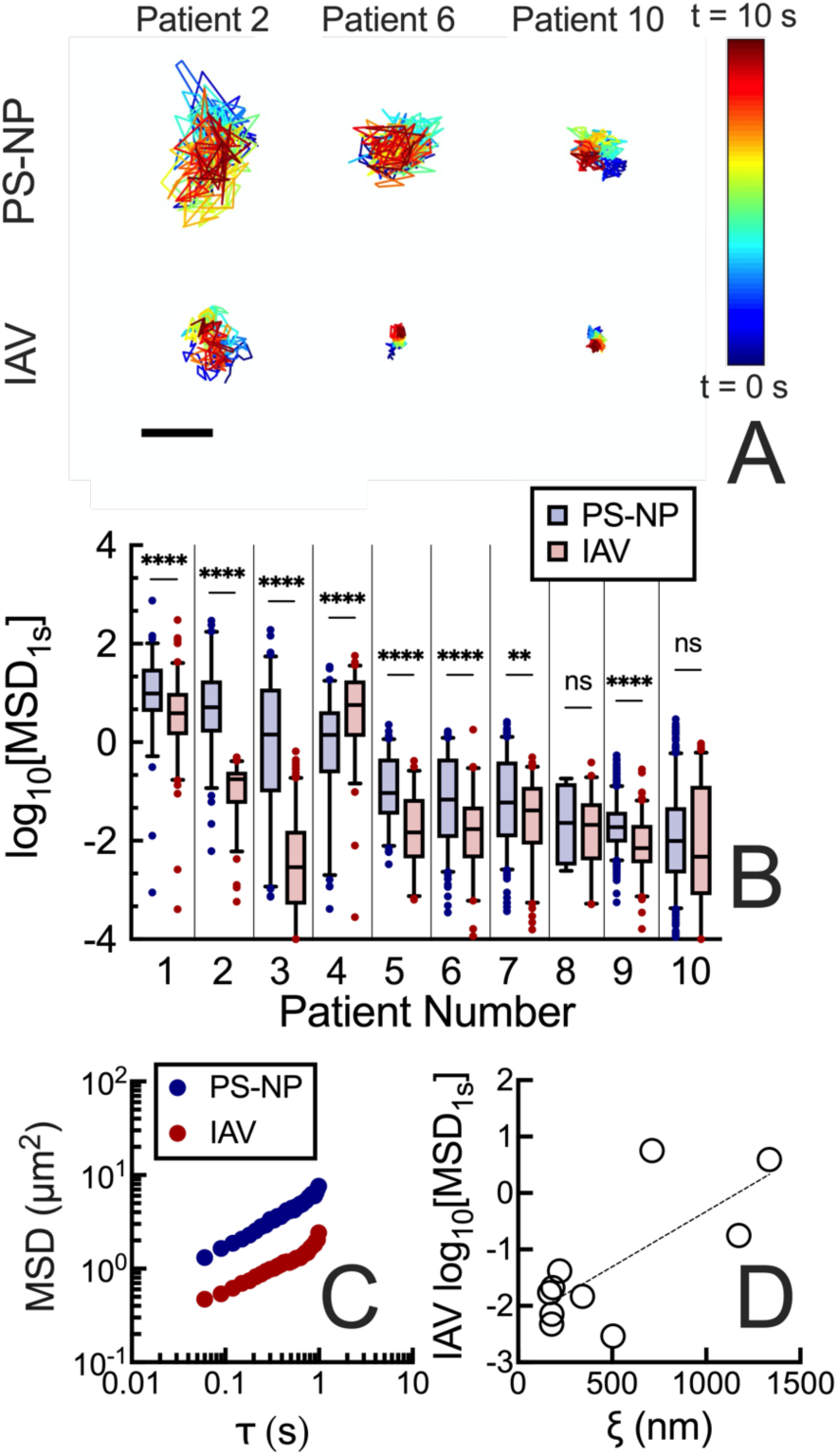
Diffusion of IAV and nanoparticles with similar diameter in human mucus. **(A**) Representative trajectories of PEG-coated 100 nm polystyrene nanoparticles (PS-NP) and IAV diffusion in mucus. Traces show 10 seconds of motion with a color scale to indicate time. The scale bar represents 0.2 µm. (**B**) Box-and-whisker plots of log based 10 of MSD at τ = 1 s (log_10_MSD) PS-NP and IAV in mucus samples collected from 10 individual patients are shown. Patients are numbered in descending order according to the median MSD of PS-NP particles in each sample. Whiskers are drawn down to the 5^th^ percentile, up to the 95^th^ percentile, and the outliers are shown as points. Data set statistically analyzed with two-tailed Mann-Whitney test: ns = not significant; p > 0.05, *p < 0.05, **p < 0.01, ***p<0.001, ****p< 0.0001. (**C**) Ensemble average MSD versus τ for IAV and PS-NP in human mucus. (**D**) Median pore size (ξ) calculated from the PS-NP MSD versus the median log_10_MSD_1s_ of IAV particles (R^2^ = 0.5468).

### Measuring the dissociation constant for IAV binding to human mucus gels

Based on our findings in **Fig. 2** showing IAV consistently diffuses more slowly than PS-NP, we reasoned IAV diffusion is influenced by specific glycan interactions mediated by HA and/or NA. In an effort to directly measure IAV interactions with glycans within the mucus gel network, we further analyzed IAV trajectories, as shown in **Fig. 2A**, using statistical mechanics. For IAV, we define an equilibrium energy of confinement (*U*) as the sum of interactions resulting from gel network viscoelasticity and hemagglutinin-mediated glycan binding. Given the comparable size of IAV and PS-NP (∼100 nm), we used PS-NP as micromechanical ‘gauges’ of network viscoelasticity as experienced by IAV. Importantly, PS-NP have been designed to exhibit negligible interactions with mucins (32). The step-by-step procedure and representative potential for PS-NP are shown in **Fig. S2**. In order to decouple the influence of physical confinement and mucin glycan binding on IAV diffusion, we subtract measured *U* for PS-NP from that measured for IAV to effectively remove the influence of local gel mechanics. With this accounted for, we calculated IAV potential energy profiles with a harmonic well potential representation (31) and a general expression was used to estimate a dissociation constant for individual IAV (31). Using this approach, the IAV trajectories were analyzed to determine the harmonic potential well depth (*U*_*M*_), spring constant (*k*_*s*_) and dissociation constant (*K*_*d*_) for binding of IAV to human mucus **(Fig. 3**). Based on our analysis, we find the mode and median for *k*_s_ of 43 and 38 pN/nm, and *K*_*d*_ of 163 and 300 mM, respectively.

**Figure 3.**
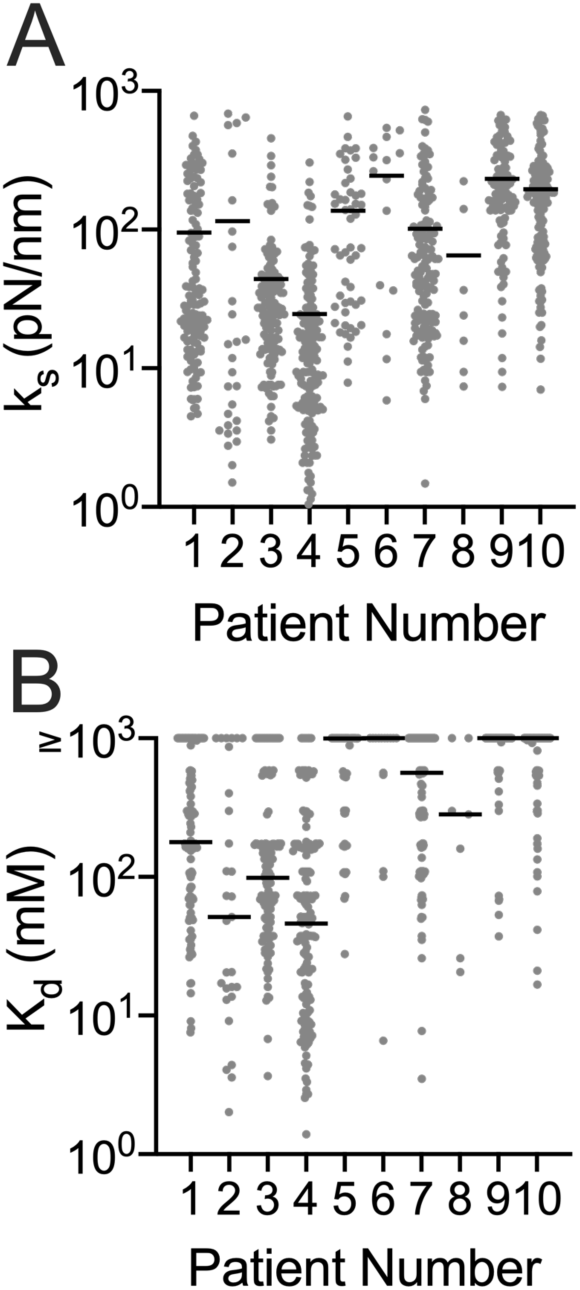
Quantifying hemagglutinin (HA) mediated IAV binding to human mucus. Scatter plots of calculated (**A**) spring constant (*k*_s_), and (**B**) dissociation constant (*K*_*d*_) for individual IAV particles. For (**B**), IAV with a calculated *K*_d_ ≥ 1000 mM are compiled with these high *K*_d_ indicative of negligible IAV-mucus interactions. Patient numbers correspond with those in Fig. 2.

### Effect of IAV neuraminidase (NA) inhibition on IAV diffusion

Given these measurable effects of glycan binding on IAV diffusion, we next examined the impact of NA activity on IAV diffusion. Prior work has shown IAV is immobilized when NA activity is inhibited given it is not be able to cleave the SA on the mucins that IAV HA binds to (13, 18). To test this, we measured IAV diffusion in human mucus in the presence of the NA inhibitor (NAI), zanamivir. Control experiments confirmed NA activity is completely abolished by NAI at a final concentration of 10 µM (**Fig. S3**). Thus, samples containing IAV and PS-NP were treated with 10 µM NAI, imaged, and subsequently analyzed with MPT. When comparing across cases, there was a significant increase in the pore size based on PS-NP diffusion in 3 of 4 samples tested after NAI treatment (**Fig. 4A**). In addition, the diffusivity of IAV significantly increased in 3 of 4 samples tested after NAI treatment (**Fig. 4B**). A small decrease in IAV diffusion was observed in 1 of 4 samples tested (patient 8), but did not significantly differ from untreated controls.

**Figure 4.**
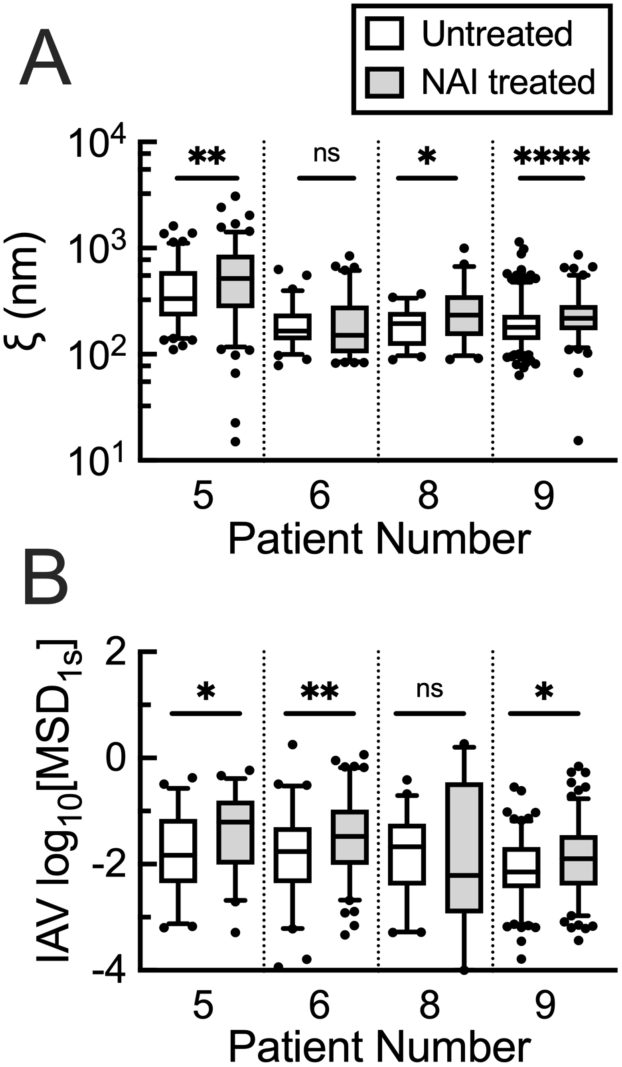
The effect of neuraminidase (NA) inhibition on IAV diffusion through human mucus. (**A**) Calculated pore size (ξ) in untreated and NA inhibitor (NAI) treated (zanamivir; 10 µM final concentration) based on PS-NP diffusion in human mucus. (**B**) Measured log_10_MSD_1s_ for IAV diffusion in untreated and NAI treated human mucus. Whiskers are drawn down to the 5^th^ percentile, up to the 95^th^ percentile, and outliers are plotted as points. Data set statistically analyzed with two-tailed Mann-Whitney test: ns = not significant; p > 0.05, *p < 0.05, **p < 0.01, ***p<0.001, ****p< 0.0001. Patient numbers correspond with those in Figs. 2 and 3.

### Disruption of mucus gel cross-linking and its impact on IAV diffusion

We next investigated the effect of cross-linking on IAV diffusion by treating samples with the mucolytic agent, N-acetyl-L-cysteine (NAC). Based on our initial data showing more rapid IAV diffusion in patient samples with greater network pore size (**Fig. 2D**), we reasoned reducing cross-linking in human mucus would result in larger network pores and allow for IAV to more readily diffuse through human mucus. To examine this, samples containing IAV and PS-NP were treated with NAC and analyzed using MPT. Once treated, both PS-NP and the IAV traveled significantly greater distances (**Fig. 5A**) leading to significant increase in measured pore size (**Fig. 5B**), and diffusion rate of the IAV particles (**Fig. 5C**).

**Figure 5.**
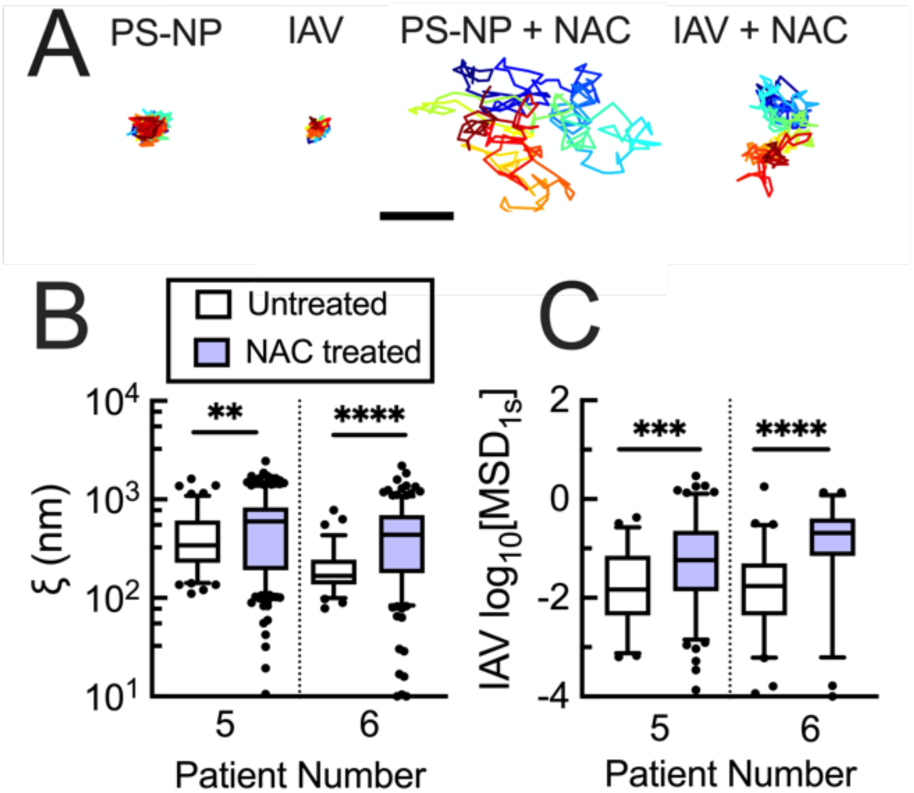
Impact of reducing disulfide bonds within mucus gels on IAV diffusion. (**A**) Representative trajectories of PS-NP and IAV diffusion in mucus with and without NAC treatment. Scale bar = 0.4 µm. (**B**) Calculated mucus gel pore size (ξ) based on PS-NP diffusion in untreated and N-acetylcysteine (NAC; 50 mM final concentration) treated mucus. (**C**) Measured log_10_MSD_1s_ for IAV in untreated and NAC-treated mucus. Data set statistically analyzed with two-tailed Mann-Whitney test: ns = not significant; p > 0.05, *p < 0.05, **p < 0.01, ***p<0.001, ****p< 0.0001. Patient numbers correspond with those in Figs. 2-4.

### Diffusion of IAV in synthetic mucus with altered microstructural properties

We then investigated the relationship between mucus cross-linking and IAV diffusion using a synthetic mucus gel model (32). We hypothesized that a systematic increase in synthetic mucus gel cross-linking and resulting tightening of gel pore structure would lead to a reduction in IAV diffusion rates. To test this, MPT experiments were performed on IAV and PS-NP in synthetic mucus of varying cross-linking densities. The total mucin concentration and associated sialic acid content was kept constant in all cases. As the crosslinker concentration increased, the pore size within synthetic mucus showed an overall decrease (**Fig. 6A**) and there were significant changes in the diffusivity of the IAV as the percentage of crosslinker increased (**Fig. 6B**). Comparison of measured pore size based on microrheology using PS-NP and the log_10_MSD values for the IAV, increased pore size corresponded with increased IAV diffusivity (**Fig. 6C**).

**Figure 6.**
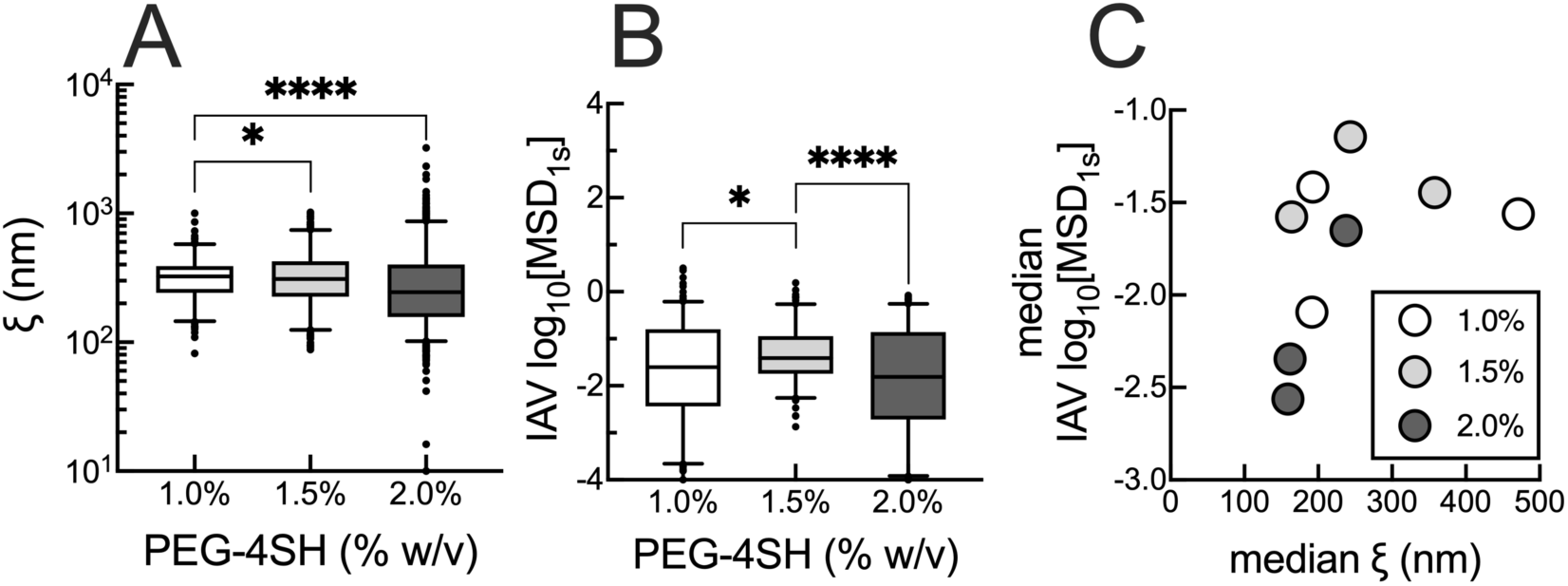
100nm PS-NP and IAV diffusion in synthetic mucus with varying crosslinking density. (**A**) Calculated pore size (ξ) based on PS-NP diffusion in synthetic mucus with varying PEG-4SH crosslinker concentration. (**B**) Measured MSD for IAV diffusion in synthetic mucus varying crosslinker concentration. (**C**) The median pore size (ξ) calculated from the PS-NP MSD compared to the median log_10_MSD value of the IAV particles in synthetic mucus with varying crosslinker concentration. Whiskers are drawn down to the 5^th^ percentile, up to the 95^th^ percentile, and the outliers are plotted as points. Data sets statistically analyzed with Kruskal-Wallis test and Dunn’s test for multiple comparisons: *p < 0.05, **p < 0.01, ***p<0.001, ****p< 0.0001.

## DISCUSSION

In this work, we provide new evidence that both gel network structure and IAV-glycan binding are involved in effective viral trapping by human mucus. Consistent with prior studies (13, 33), IAV displayed lower mobility in human mucus than muco-inert PS-NP with comparable diameter (∼100 nm) presumably as a result of IAV binding to the mucus gel network. We developed new analyses to decouple the impact of network structure and glycan binding on IAV diffusion behavior in human mucus. *K*_d_ determined for individual IAV virions in all human mucus samples tested were within a mM range, consistent for protein-sugar mediated binding (34, 35). These relatively weak IAV-mucus interactions alone are unlikely to provide sufficient means to immobilize IAV, but likely contribute to the slowing of IAV diffusion as compared to the non-mucoadhesive PS-NP. Further, we have demonstrated that a smaller network size, as detected by PS-NP, substantially decreases IAV mobility in human mucus and a synthetic mucus model with systematically varied crosslink density. We also found disrupting mucus crosslinking with the reducing agent NAC enhanced IAV diffusion through mucus. In addition, we considered the possibility that IAV NA could degrade mucus and disrupt its structure to promote penetration, similar to the mechanism used by bacterial pathogens (36). However, we observed an increase in PS-NP diffusion when incubated with IAV in only 1 of 3 patient samples tested (**Fig. S4**).

Several reports on IAV diffusion through mucus suggest interactions with mucin-associated glycans contribute to its transport behavior (13, 18, 37). For example, it has been shown H1N1 and H3N2 IAV adhere to human lung tissue sections and salivary mucus in a sialic acid-dependent manner (16). In addition, previous work has shown NA activity of a swine IAV facilitates its transit through porcine lung mucus (13). Conversely, we have found in our studies NA inactivation does not reduce IAV diffusion through mucus. The unexpected increase in network size and IAV mobility observed after NAI treatment (**Fig. 4**) may be explained by either direct effects of NAI on mucus structure or differences between aliquots from individual donors. Still, our results would suggest NA is a minor factor in IAV diffusivity in mucus. Our findings that NA is not required for IAV mobility are consistent with a recent report on IAV diffusion in human airway mucus, using the same PR8 IAV strain and human mucus from ETs (17). Furthermore, it is likely IAV HA binding of mucin glycans is weak and reversible in the presence or absence of NA activity. Given the differences in the properties of mucus from different species and mucosal tissues (38), we envision these results using a human IAV and patient-derived lung mucus provide a representative picture of their dynamics in the airway microenvironment.

We have also developed a new analysis capable of extracting effective binding constants for IAV within the mucus gel network. Unlike standard approaches using glycan binding arrays with purified viral components (39, 40), we are able to directly determine disassociation constants for intact IAV within native mucus gels. Importantly, our analysis allows us to account for the physical pore structure of mucus gels with analysis of PS-NP confinement energies. The calculated *k*_*s*_ for IAV-mucus binding, with median value of 43 pN/nm, is on the order of forces required to disrupt HA-sialic acid bonds (10-25 pN (41, 42)). Calculated *K*_d_ values varied widely between patients over a mM range, but are generally consistent with short-lived, reversible HA-sialic acid bonds with reported *K*_d_ ranging from 1-20 mM (18). We attribute the large range in measured *K*_d_ to differences in the properties of mucus collected rather than functional differences in individual IAV. There is likely wide variation in the glycans present in human mucus from different donors and local differences in the spatial arrangement of glycans within distinct regions of mucus gels. These estimates of *K*_d_ would suggest IAV HA is limited to monovalent interactions rather than multi-valent interactions within the mucus network. The affinity of IAV HA towards mucus may also be reduced by the diversity of terminal glycans (e.g. sialic acid, sulfate, fucose) expressed in airway mucus (43). Interestingly, we observed the lowest glycan affinity in IAV diffusion in patient samples 5-10 (**Fig. 4B**) with dense mucus gel networks (**Fig. 2B**). Relevant to these observations, a recent study demonstrated the inhibitory effect of high mucin density on IAV binding to supported lipid bilayers engineered with surface-tethered synthetic mucin mimetics (44). Steric inhibition of HA-sialic acid binding within dense and/or highly crosslinked mucus may explain the estimates of *K*_d_ ≥1000 mM, indicative of negligible HA-mediated glycan binding.

Our data supports the rationale that enhanced concentration and crosslinking of mucus gels can help to protect the airway epithelium from IAV and potentially other respiratory viruses. The inflammatory response to IAV infection is known to increase reactive oxygen species (ROS) production and cytokine expression resulting in acute lung injury (45). In response to ROS and oxidative stress, the epidermal growth factor (EGFR) pathway, responsible for mucin hypersecretion, becomes activated leading to increased mucus concentration in the airway lumen (46, 47). In addition, prior studies have found ROS and oxidative stress can increase the elasticity of airway mucus (43), attributed to the oxidation of mucin cysteine domains leading to formation of mucin-mucin disulfide crosslinks (48). Increased mucin concentration and mucin-mucin crosslinking due to oxidative stress might be a beneficial part of the inflammatory response towards IAV infection to hinder virus penetration through the mucus barrier. These potential benefits, however, may be undermined by compromised airway clearance in mucus gels with abnormal viscous and elastic properties (10).

In summary, we have shown IAV particle diffusion through mucus gels is limited by both the structural properties of mucus and mucin glycan recognition by IAV HA. The influence of gel network structure observed in our work bares similarity to past studies on particle diffusion through mucus and other biological matrices (49–53). IAV was found to be weakly adherent to human mucus which likely improves its ability to navigate to the underlying airway epithelium. Our results yield new and important insight into influenza A infection of the airway. All of the studies reported here were carried out using the PR8 strain of IAV. To determine the broader relevance of these findings, we plan to include alternative strains in future work with different sialic acid preference and more contemporary IAV strains where neutralizing antibody-mediated entrapment may influence IAV-mucus interactions (33). We also note PR8 and other lab adapted IAV strains produce primarily spherical virions (54). In natural infections, influenza virions can vary in morphology, producing a mixture of spherical and filamentous virions (54). Recent work has also shown the impact of IAV particle morphology on transport in mucus which will be important to consider in subsequent studies (55).

## CONCLUSION

In this work, we determined that IAV mobility can be limited by both the structural and biochemical features of the mucus gel network. Unlike previous reports, we found NA on the IAV capsid is not required to enable penetration through mucus gel networks. As might be expected, mucus gels with pore sizes approaching the diameter of IAV are capable of physically entrapping viral particles. Our measurements suggest IAV binding to mucus is weak and reversible due to low affinity interactions between HA and sialic acid. Steric hindrance within the mucus gel network may also influence accessibility of sialic acid receptors to HA binding. Our results also provide a framework for direct measurement of viral particle association to mucus gels which may prove useful for future studies on respiratory viruses by our group and others in the field.

## AUTHOR CONTRIBUTIONS

L.K., conceived, designed, and performed the research and led development of data analysis to determine virus binding affinity. E.I. designed and performed experiments and produced all viruses used for this study. S.B. and D.S. helped with experiments including tracking experiments and synthetic mucus formulation. M.A.S. and G.A.D. conceived and designed experiments. L.K. and G.A.D. wrote the article. All authors reviewed and edited the article.

## ACKNOWLEDGEMENTS

This project was funded by NIH R21 AI142050 (to M.A.S. & G.A.D), Burroughs Welcome Fund CASI (to G.A.D.), the Parker B. Francis Fellowship Program (to M.A.S.), and the American Lung Association (to G.A.D.). We would also like to thank Dr. Irina Timofte and Anu Varghese at UMMC for collecting the human mucus samples used in these studies.

