## Supplemental Information for "Influenza A virus diffusion through mucus gel networks"

SUPPLEMENTAL FIGURES

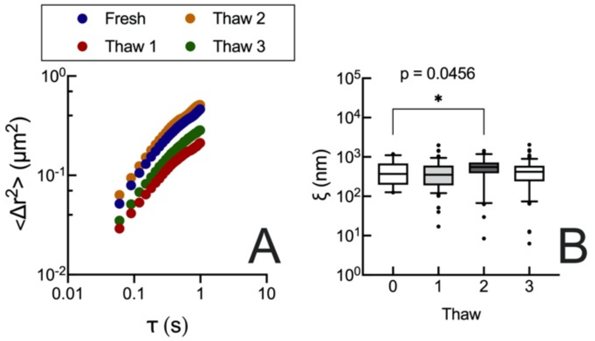


**Figure S1. Human mucus sample with varying freeze-thaws**
(**A**) MSD of particles dispersed in a fresh HM sample and after 1, 2, and 3 freeze-thaw cycles. (**B**) Calculated pore size ($\xi$) for individual particles at $\tau = 1$ s. Whiskers are drawn down to the 5$^{th}$ percentile, up to the 95$^{th}$ percentile, and the outliers are plotted as points. Data sets statistically analyzed with two-tailed Mann-Whitney test: *$P < 0.05$, **$P < 0.01$, ***$P<0.001$, ****$P< 0.0001$.

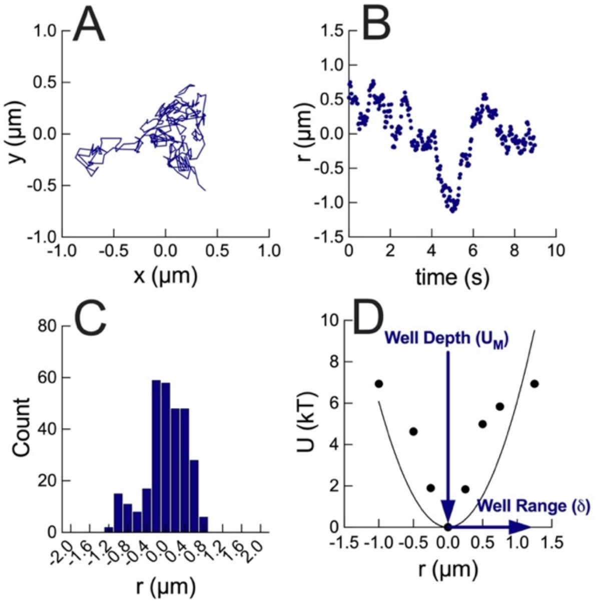


**Figure S2. Displacement and equilibrium energy of confinement of individual PS-NP**
(**A**) PS-NP trajectory centered around average x and y value. (**B**) Particle displacement (*r*) from the trajectory center over time. (**C**) Distribution of particle displacement over time. (**D**) Equilibrium energy of confinement (*U*) vs displacement (*r*) with well depth and well range labeled.

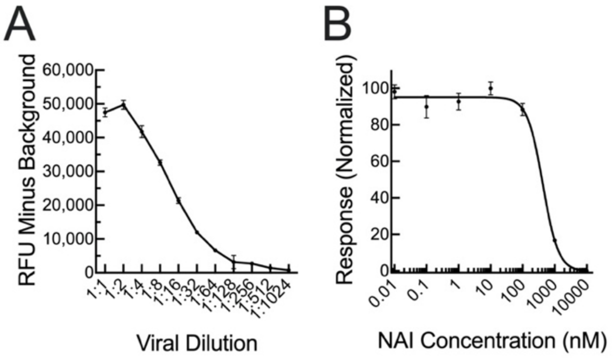


**Figure S3. Neuraminidase activity and inhibition assay for unlabeled IAV**
(**A**) Neuraminidase activity at different viral dilutions of unlabeled IAV. (**B**) Neuraminidase activity of unlabeled IAV (1:16 dilution) in varying concentrations of neuraminidase inhibitor Zanamivir.

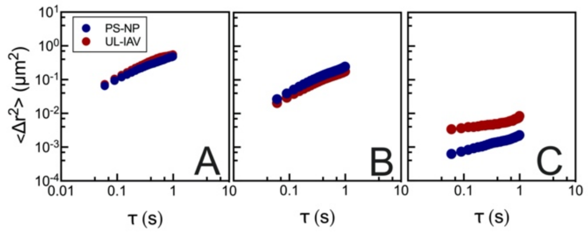


**Figure S4: 100nm PS-NP and unlabeled IAV human mucus samples**
(**A**) MSD of particles dispersed in human mucus samples. (**B**) Measured $log_{10}$MSD for individual particles at $\tau = 1$ s across all samples. (**C**) Calculated pore size ($\xi$) for PS-NP at $\tau = 1$ s across all samples. Whiskers are drawn down to the $5^{th}$ percentile, up to the $95^{th}$ percentile, and the outliers are plotted as points. Data sets statistically analyzed with two-tailed Mann-Whitney test: *$P < 0.05$, **$P < 0.01$, ***$P<0.001$, ****$P< 0.0001$.
